# The ciliary membrane of polarized epithelial cells stems from a midbody remnant-associated membrane patch with condensed nanodomains

**DOI:** 10.1101/667642

**Authors:** Miguel Bernabé-Rubio, Minerva Bosch-Fortea, Esther García, Jorge Bernardino de la Serna, Miguel A. Alonso

## Abstract

The primary cilium is a specialized plasma membrane protrusion that harbors receptors involved in important signaling pathways. Despite its central role in regulating cellular behavior, the biogenesis of the primary cilium is not fully understood. In fact, the source of the ciliary membrane remains a mystery in cell types that assemble their primary cilium entirely at the cell surface, such as polarized renal epithelial cells. After cytokinesis, the remnant of the midbody of these cells moves to the center of the apical surface, where it licenses the centrosome for ciliogenesis through an unidentified mechanism. Here, to investigate the origin of the ciliary membrane and the role of the midbody remnant, we analyzed membrane compaction and lipid dynamics at the microscale and nanoscale in living renal epithelial MDCK cells. We found that a specialized patch made of condensed membranes with restricted lipid lateral mobility surrounds the midbody remnant. This patch accompanies the remnant on its journey towards the centrosome and, once the two structures have met, the remnant delivers part of membranes of the patch to build the ciliary membrane. In this way, we have determined the origin of the ciliary membrane and the contribution of the midbody remnant to primary cilium formation in cells whose primary cilium is assembled at the plasma membrane.

## Introduction

The primary cilium consists of a single membrane protrusion containing a microtubule-based scaffold, termed the axoneme, which is nucleated from the older of the two centrioles in the centrosome. The ciliary membrane is continuous with the plasma membrane but differs from it in structure and composition. The ciliary membrane harbors a large variety of receptors for cell signaling, including those for soluble factors involved in cell growth, migration, development and differentiation, and G-protein-coupled receptors (Ishikawa and Marshall, 2011; Gerdes, et al., 2009; Singla and Reiter, 2006). Abnormalities in ciliary membrane functioning result in a growing list of human developmental and degenerative disorders that simultaneously affect nearly every major organ, especially the kidney (Braun and Hildebrandt, 2017; Reiter and Leroux, 2017).

The origin of the ciliary membrane depends on the route of primary cilium formation used (Bernabé-Rubio, et al., 2017; Sorokin, 1968). In cells in which the process of primary cilium formation starts intracellularly, as occurs in fibroblasts, the ciliary membrane derives from a vesicle that progressively expands at the distal part of the mother centriole. The vesicle then gradually deforms by elongation of an incipient axoneme in such a way that, upon its exocytosis, the membrane on the side of the vesicle facing the axoneme becomes the ciliary membrane (Sorokin, 1962). In contrast, in cells, such as renal polarized epithelia, in which the process of primary cilium formation takes place by an alternative route occurring entirely at the cell surface, the source of the ciliary membrane remains a mystery. Given its relevance, understanding how these cells manufacture the ciliary membrane is a matter of fundamental biological importance and can broaden our understanding of the primary cilium-associated diseases.

The intercellular bridge formed during cytokinesis, also known as the midbody, contains in the middle an electrodense structure called the Flemming body that is either released after abscission to the extracellular space or inherited as a midbody remnant (MBR) by one of the daughter cells (Chen, et al., 2012; Green, et al., 2012). The MBR serves a role as a polarity cue in cell processes such as neurite outgrowth (Pollarolo, et al., 2011) or epithelial lumen formation (Li, et al., 2014). Stem cells and cancer cells enriched in midbody remnants exhibit increased reprogramming efficiency and *in vitro* tumorigenicity, respectively (Ettinger, et al., 2011; Kuo, et al., 2011). Although the exact role of the MBR in all these processes is unknown, its importance in cellular physiology and cell fate determination is increasingly apparent (Dionne, et al., 2015; Chen, et al., 2012). In polarized Madin-Darby canine kidney (MDCK) cells, which are a paradigm of renal tubular epithelial cells (Rodriguez-Boulan, et al., 2005), the MBR membrane remains continuous with the plasma membrane through a thin stalk derived from the unresolved side of the intercellular bridge. The MBR then moves along the apical membrane to meet the centrosome and thereby enable primary cilium assembly (Bernabe-Rubio, et al., 2016). However, what the MBR does to license the centrosome for primary cilium assembly is not known. Since primary cilia contain condensed membranes (Vieira, et al., 2006), in this study we have investigated the possibility that the ciliary membrane of cells following the alternative route arises from specialized membranes provided by the MBR. By monitoring condensed membranes in live cells using state-of-the-art super-resolution and fluorescence correlation spectroscopy-based techniques, we have revealed the origin of the ciliary membrane and the role of the MBR in primary ciliogenesis in polarized epithelial cells.

## Results and Discussion

### The midbody remnant is enriched in condensed membranes

Membrane condensation can be measured with polarity sensitive dyes, such as Laurdan (6-lauryl-2-dimethylamino-napthalene) and di-4-ANEPPDHQ, that report lipid packing in model and cell membranes (Owen, et al., 2011). The fluorescent membrane probe Laurdan undergoes a 50-nm shift in its peak emission wavelength from 440 nm in highly ordered condensed membranes to 490 nm in disordered fluid membranes. A normalized ratio of the intensity at the two emission regions, 420-460 nm and 470-510 nm, which is given by the generalized polarization (GP) index, whose values range from −1 (most fluid) to +1 (most condensed), provides a relative measure of lipid order in cell membranes (Gaus, et al., 2003; Parasassi, et al., 1994). It is of note that lipid organization is best measured in live cells because fixation can induce artifacts (Tanaka, et al., 2010). Therefore, to analyze membrane condensation at the MBR zone, live renal epithelial MDCK cells stably expressing mCherry-MKLP1, which labels the MBR’s interior, were stained with Laurdan. GP values were calculated for each pixel and then used to generate a color-scaled image to visualize the degree of membrane condensation. An apical ring-shaped patch of condensed membranes, hereafter referred to as a remnant-associated membrane patch (RAMP), was found surrounding MBRs of MDCK (Figs. 1A, B) and of the inner medullary collecting duct 3 (IMCD3) renal cell line (Fig. S1A), in which approximately 90% of cells assemble the primary cilium at the cell surface (Molla-Herman, et al., 2010).

**Figure 1.**
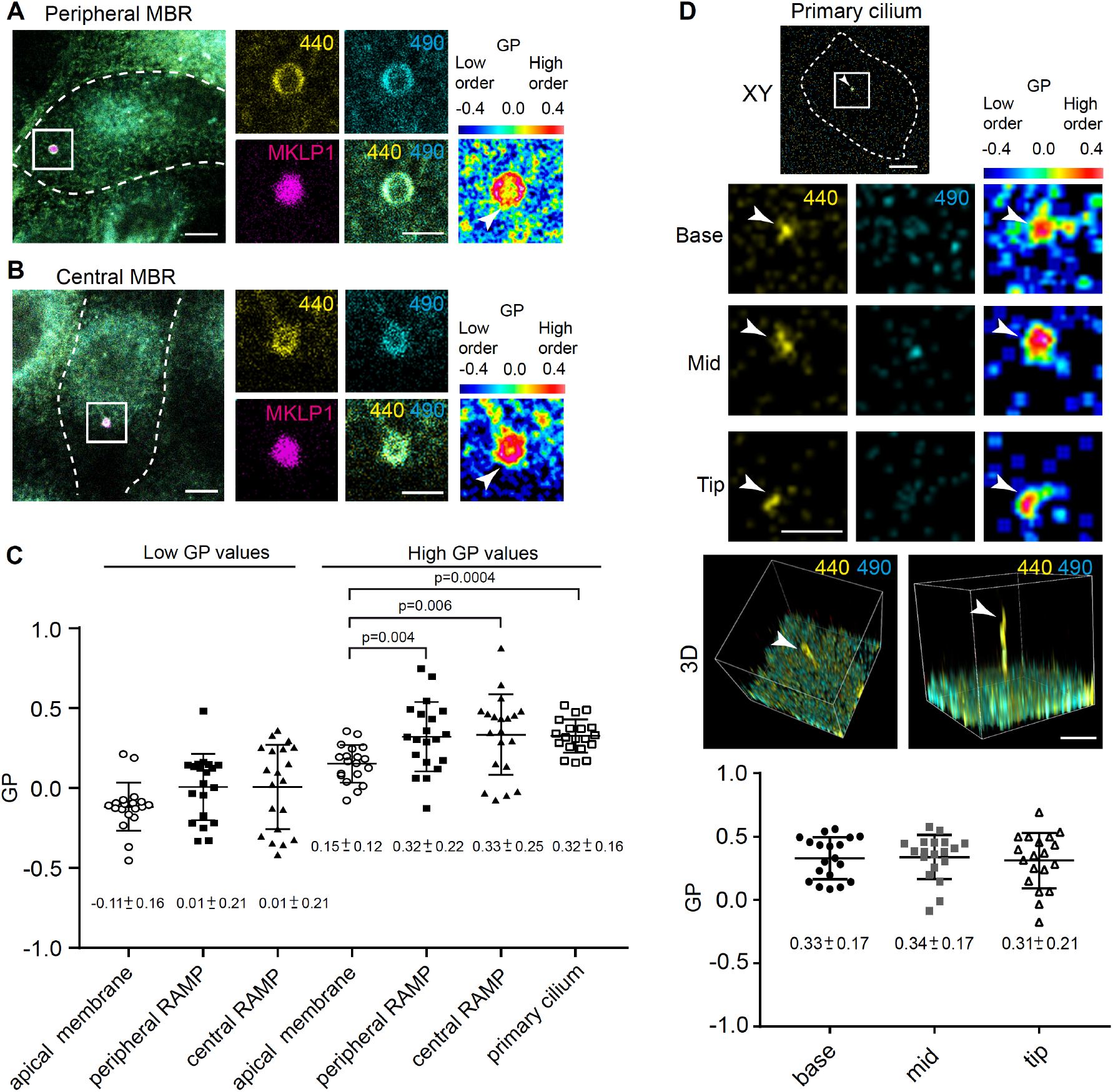
A patch of condensed membranes is associated with the MBR. **(A, B)** MDCK cells stably expressing mCherry-MKLP1 were stained with Laurdan. Representative XY projections of cells with either a peripheral (A) or a central MBR (B) are shown. The enlargements of the boxed zone show the emission at λ440 and λ490 nm in the MBR area, the corresponding merge and the localization of mCherry-MKLP1. Arrowheads indicate the MBR. **(C)** GP values corresponding to different structures at the apical membrane. MBR zones were delimited by rings; identical rings were used to delimit membrane regions chosen at random. For the cilium, we show the mean of the GP value found in cross-sections at the base, middle and tip. 19-21 structures were examined in each case. **(D)** The enlargements show representative XY images of cross-sections at the base, middle and tip of a primary cilium of the emission at λ440 and λ490 nm, and the corresponding GP images of cells stained with Laurdan. 3D projections of a representative primary cilium are shown below. Arrowheads indicate the primary cilium. The bottom histogram shows the mean GP value at the base, middle and tip cross-sections of primary cilia (19 cilia examined). Dashed lines indicate the cell contour. Mean values ± SD are presented in C and D. At least three independent experiments were performed in C and D. Scale bars, 5 µm for panoramic views and 2 µm for enlargements. Colored scales are shown next to each set of GP images.

To analyze quantitatively the lipid organization of RAMPs, the distribution of GP values obtained in MDCK cells was fitted to two Gaussian populations (Fig. S1B). We found a more fluid membrane population with GP values of 0.01 ± 0.21, and a more condensed membrane population with GP values of 0.32 ± 0.22 at peripheral RAMPs (Fig. 1C). The coverage, which represents the proportion of condensed domains, was 55.84 ± 21.37% of the total membranes in the peripheral RAMPs. RAMPs at central MBRs showed similar membrane condensation (GP = 0.33 ± 0.25, Fig. 1C) and coverage (57.08 ± 25.86%). The degree of condensation at RAMPs was significantly higher than that of the apical plasma membrane (Fig. 1C). Similar GP values were obtained from IMCD3 cells (Fig. S1C).

Analysis of live MDCK and IMCD3 cells revealed that the ciliary membrane almost exclusively comprises condensed membranes (coverage = 82.91 ± 22.09%) and is similar in its degree of condensation (GP = 0.34 ± 0.17) to that of RAMPs (Figs. 1C, D, Figs. S1C, D). The GP values along the primary cilium (base, middle and tip cross-sections) were within the same range (Fig. 1D), indicating membrane continuity with similar lipid lateral packing from the base to the ciliary tip. In conclusion, the results in Figure 1 illustrate a similarity in the degree of membrane condensation in peripheral and central MBRs and in the ciliary membrane, which in turn are very different from that of the plasma membrane.

### The condensed membranes at the RAMP have limited lipid lateral mobility and are sensitive to cholesterol depletion

Condensed membranes in cells, sometimes referred to as rafts or raft-like membranes (Sezgin, et al., 2017; Farnoud, et al., 2015), were proposed to serve as functional protein signaling platforms (Lingwood and Simons, 2010; Simons and Gerl, 2010). Since condensed membranes contain a high proportion of cholesterol (Simons and Sampaio, 2011), we examined the effect of depleting cholesterol in live cells using either cholesterol oxidase or methyl-β-cyclodextrin. In both cases, we observed that the percentage of cells with a RAMP greatly diminished (Figs. S1E, F). This result supports that RAMPs consist of condensed membranes.

A critical property of condensed yet fluid membranes compared with more rigid membranes is their ability to allow molecular lateral mobility, although in a more restricted manner than in fluid membranes (Bernardino de la Serna, et al., 2016). To examine lipid lateral diffusion we used raster imaging correlation spectroscopy (RICS), which takes advantage of the correlation of the fluorescence fluctuations over time to yield diffusion coefficient values of molecules (Garcia and Bernardino de la Serna, 2018; Digman, et al., 2005). We labeled cells with di-4-ANEPPDHQ, which gives better videomicroscopy perfomance than with Laurdan due to its higher signal-to-noise ratio and photostability (Owen, et al., 2011), to record membrane compaction and lipid lateral mobility simultaneously. The emission of di-4-ANEPPDHQ was recorded at 500-580 nm and 620-750 nm with peaks at 560 and 620 nm, respectively. 3D reconstruction of cells expressing mCherry-MKLP1 shows that the RAMP surrounds the MBR (Fig. 2A). The analysis of lipid diffusion at different optical planes along the z-axis of the RAMP revealed that the diffusion coefficient of di-4-ANEPPDHQ was much lower at RAMPs than at the apical membrane (Fig. 2A, B). The analysis of membrane compaction at the RAMP with di-4-ANEPPDHQ (Fig. 2B) confirms the result obtained with Laurdan (Fig. 1C) and further suggests that the degree of membrane condensation at RAMPs is much higher than at the apical membrane. It is also of note that the diffusion coefficients at different planes of the RAMP were in the same range (Fig. 2B), indicating the continuity of collective motion and condensation of lipids in the RAMP.

**Figure 2.**
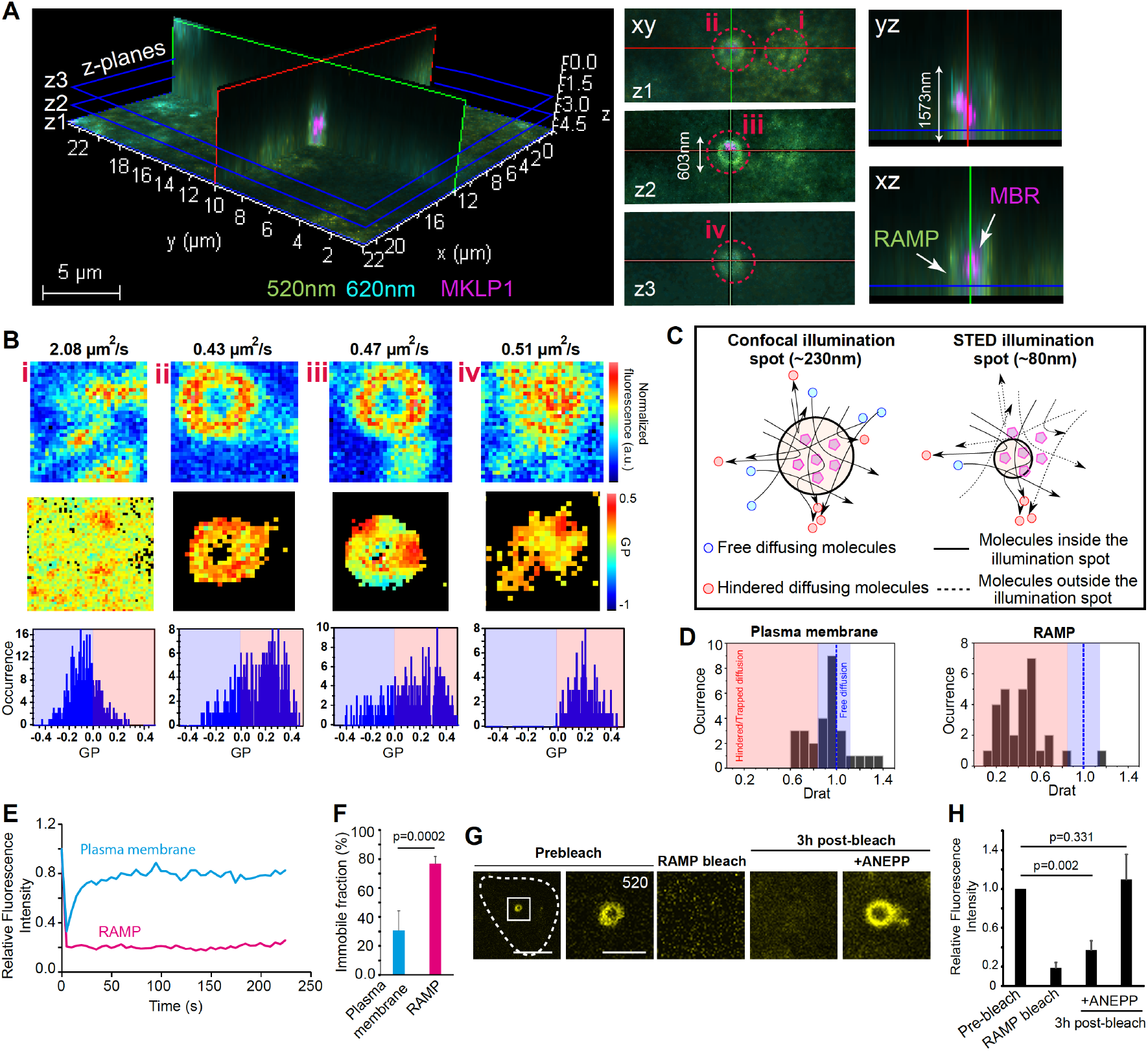
The RAMP is a specialized membrane domain of compact membranes. **(A)** XY, YZ and XZ projections of an MBR at the apical membrane of MDCK cells. Cells stably expressing mCherry-MKLP1 (magenta) were labeled with di-4-ANEPPDHQ. The emission at λ560 and λ620 nm are shown in green and cyan, respectively. **(B)** Optical cross-sections z1-3 from panel A show apical membrane (inset i), and proximal (ii), middle (iii) and distal (iv) planes of a central RAMP. Lipid diffusion and membrane condensation analyses were performed. Panels from top to bottom: diffusion coefficient and diffusion map obtained from RICS analysis, GP map, and histograms of GP frequency (low GP and high GP values in light-blue and light-red backgrounds, respectively). **(C)** Schematic of confocal and STED imaging volumes shows likelihood of detecting hindered diffusion of molecules in a smaller imaging spot size. **(D)** The histograms show the Drat ratio at the plasma membrane and at the RAMP. Three independent experiments; 27-28 regions of interest were analyzed. **(E)** MDCK cells were labeled with di-4-ANEPPDHQ and subjected to FRAP for 220 s in the RAMP and in plasma membrane. A representative example of the kinetics of recovery of di-4-ANEPPDHQ fluorescence after photobleaching is shown. **(F)** The histogram depicts the percentage of the immobile fraction at RAMPs and at the plasma membrane (10 cells were analyzed). **(G, H)** 3 h after RAMP photobleaching, the cells were re-stained with di-4-ANEPPDHQ to visualize the bleached RAMP (G). (H) The histogram shows the relative fluorescent intensity at the indicated steps of the process. Mean values ± SEM are represented in (F, H). Enlargements of the boxed region are shown. Scale bars, 5 µm for panoramic views and 2 µm for enlargements.

The length and time scale resolution of RICS can be diminished by reducing the focal observation volume. Stimulated emission depletion (STED) nanoscopy is capable of measuring subdiffraction-limited molecular events by decreasing the point spread function volume or illumination volume of a time-lapse acquisition. Thus, by combining STED with RICS (STED-RICS microscopy), we can assess the spatial molecular localization and the lipid lateral mobility at the nanoscale resolution (Hedde, et al., 2013). Based on a recently reported technique in which alternating line scanning was used instead of RICS (Schneider, et al., 2018), we applied the same principle to RICS and simultaneously acquired confocal and STED images on a line-by-line basis to examine lipid nanodomain dynamics at RAMPs (Fig. 2C). By resolving the diffusion of the molecules simultaneously in diffraction-limited (D) and super-resolution (D_STED_) modes, different mobility modes can be discerned (Schneider et al., 2018). For instance, the ratio between the quantified diffusion values (D_STED_/D), hereafter referred to as Drat, allows discrimination among free diffusion, hindered or trapped diffusion, and active diffusion. If Drat = 1, lipids move randomly at the plasma membrane. However, if Drat < 1, as occurs in RAMPs (Fig. 2D), lipids are trapped because their diffusion is hindered by the presence of highly condensed raft-like nanodomains. Therefore, this analysis indicates that, similar to the way lipids would move in raft-like enriched membranes immersed in a more fluid environment, molecular lipid mobility is hindered at RAMPs whereas, by contrast, such restriction was not detected at the plasma membrane.

Complementary analyses using fluorescence recovery after photobleaching (FRAP) showed that the turnover of lipids at RAMPs was much lower than that of the plasma membrane (Figs. 2E). In addition, the percentage of the immobile lipid fraction present at RAMPs was significantly higher than that of the plasma membrane (Fig. 2F), confirming that RAMPs are specialized lipid domains. The level of lipid packing/condensation found in RAMPs was such that the fluorescent signal at the RAMP had not recovered 3 h after having been photobleached. Only after re-staining the cells with new di-4-ANEPPDHQ did the RAMP recover its fluorescence, indicating that the RAMP lipids are not replaced by lipids diffusing from the plasma membrane or from intracellular vesicles once the RAMP has formed (Fig. 2G, H). Together, these results support the existence of a patch of condensed membranes at the MBR zone with low lipid lateral mobility. The finding that RAMPs are specialized membranes is consistent with previous studies showing that cells specifically regulate the localization of lipids to the midbody (Atilla-Gokcumen, et al., 2014), and with lipidomic analysis indicating that the midbody has a different lipid composition from that of most cellular membranes (Arai, et al., 2015).

### The midbody remnant delivers condensed membranes to the centrosome

Whereas the MBR is lost from subconfluent cultures of MDCK cells (Elia, et al., 2011), it is maintained at the cell periphery of cells grown at high cell density as a membrane protrusion that is continuous with the apical surface through a thin membranous tether (Bernabe-Rubio, et al., 2016). To determine when the MBR acquires the RAMP, we monitored the progression of cytokinesis by time-lapse microscopy of Laurdan-stained cells expressing mCherry-tubulin to visualize the microtubules in the intercellular bridge and in the MBR. The presence of the RAMP was evident soon after separation of the sister cells (Fig. S2A). The RAMP then accompanied the MBR in its movement to the center of the apical membrane, as observed in MDCK cells expressing mCherry-MKLP1 (Fig. S2B).

Before the MBR and the centrosome met, the centrosome zone lacked condensed membranes (Fig. 3A). It is of particular note that once the RAMP reached a central position, a second patch of condensed membranes started to appear adjacent to the MBR (Fig. 3B). The RAMP progressively fed the emerging patch with membranes and both patches eventually separated in such a way that the remainder of the RAMP continued to be associated with the MBR, whereas the new patch localized to the plasma membrane zone above the centrosome (Fig. 3C and Movie 1). To distinguish the two membrane patches, we referred to the newly formed patch as the centrosome-associated membrane patch (CAMP). After separation of the patches, the MBR alongside the RAMP eventually disappeared, as determined by the loss of mCherry-MKLP1 labeling (Fig. S2C). Physical removal of peripheral MBRs from the cell surface, which impairs primary cilium formation (Bernabe-Rubio, et al., 2016), greatly reduced the percentage of cells with a CAMP, confirming that the CAMP originates from the RAMP (Fig. 3D and Fig. S3D).

**Figure 3.**
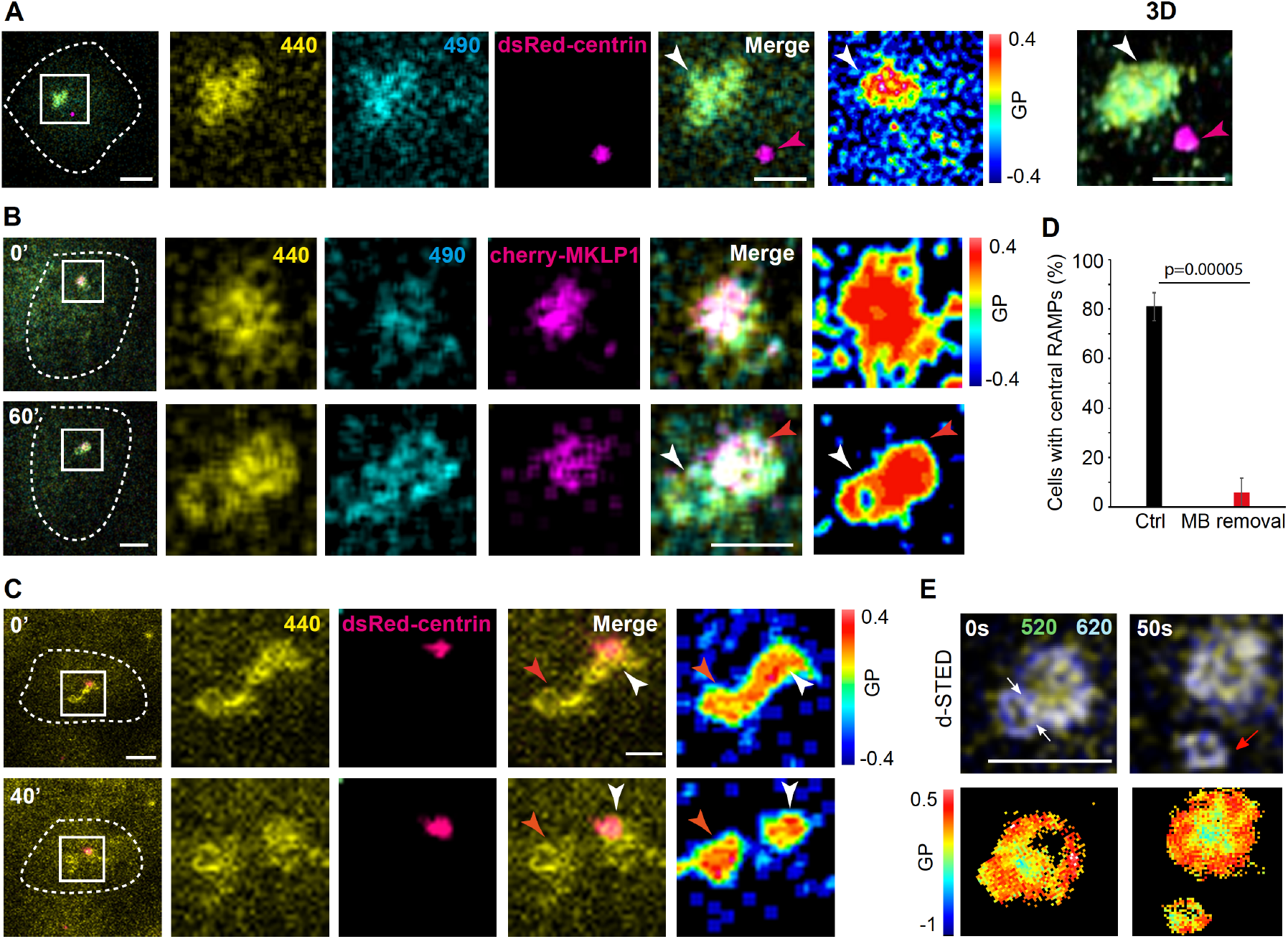
The MBR delivers a membrane patch to the centrosome. **(A)** Cells expressing dsRed-centrin were stained with Laurdan. The enlargements of the boxed zone show the emission at λ440 and λ490 nm of a RAMP close to the centrosome, the corresponding merge and GP images, and the 3D reconstruction. Note the absence of Laurdan fluorescence at the centrosome zone when the RAMP is near the centrosome but has not yet reached it. **(B, C)** Cells were stained with Laurdan and analyzed by videomicroscopy. The images show the Laurdan emission at λ440 and λ490 nm of a central RAMP close, and the position of the MBR and the centrosome, as visualized with mCherry-MKLP1 (B) and dsRed-centrin (C), respectively, and the corresponding merge and GP images. Note in (B) that the RAMP starts remodeling to form an adjacent patch, which is called a CAMP. The white and red arrowheads point to the CAMP and the remaining RAMP membranes, respectively. In (C) the original RAMP separates into two patches, the new one being associated with the centrosome. **(D)** The MBR of MDCK cells stained with Laurdan was removed by aspiration. As a control (Ctrl), the same procedure was applied in a zone of the plasma membrane distant from the MBR. The percentage of cells showing a CAMP was quantified 24 h later. Results are summarized as the mean ± SEM of three independent experiments (*n* = 35 control cells and 30 cells whose MBR had been removed). **(E)** Cells stained with di-4-ANEPPDHQ were subjected to super-resolution videomicroscopy. Top panels, merged images of deconvolved STED of the emission at λ560 (in yellow) and λ620 nm (in blue); bottom panels, corresponding GP maps. Note the concentration of highly compact membranes at the zone of CAMP emergence (white arrows). The red arrow indicates a nascent CAMP. The dashed lines indicate the cell contour. Scale bars, 5 µm for panoramic views and 2 µm for enlargements.

Super-resolution videomicroscopic analyses showed that condensed membranes at the RAMP remodel and concentrate at the site of CAMP formation (Fig. 3E) and at the interface between the nascent CAMP and the RAMP (Fig. S2E). This arrangement could result in an increase of membrane tension and, consequently, in membrane rupture and separation of the two patches. Consistent with the accumulation of condensed membranes at the zone of CAMP formation, we found that this region had a diffusion coefficient lower than of the rest of the RAMP (Fig. S3A). In summary, Fig. 3 indicates that the RAMP moves together with the MBR to the center of the cell and gives rise to a new membrane patch that is transferred to the centrosome.

### The ciliary membrane builds from CAMP membranes

The presence of condensed membranes at the ciliary membrane is consistent with previous reports in the primary cilium of MDCK cells (Vieira, et al., 2006) and in the flagellum of *T. brucei* ((Tyler, et al., 2009). The source of these specialized membranes in cells, such as epithelial MDCK cells, that assemble the primary cilium entirely at the plasma membrane is unknown. We reported that the MBR moves along the apical membrane to meet the centrosome and somehow enables it to form a primary cilium (Bernabe-Rubio, et al., 2016). To investigate whether the MBR licenses the centrosome by providing CAMP membranes to build the ciliary membrane, we analyzed by time-lapse microscopy the process of primary cilium formation in Laurdan-stained cells expressing dsRed-centrin. We observed that after the encounter between the MBR and the centrosome, the axoneme began to emerge surrounded by the ciliary membrane that, as shown before (Fig. 1D), is comprised of condensed membranes (Fig. 4A and Movie 2). This result is consistent with a previous report showing that proper membrane condensation at the centrosome zone is necessary for efficient primary cilium assembly (Reales, et al., 2015). It is of note that the CAMP decreased in size as the cilium elongated, as seen in 3D reconstructions (Fig. 4A and Movie 2). Since no other condensed membranes were detected near the growing cilium, these observations suggest that CAMP membranes are used for the assembly of the ciliary membrane.

**Figure 4.**
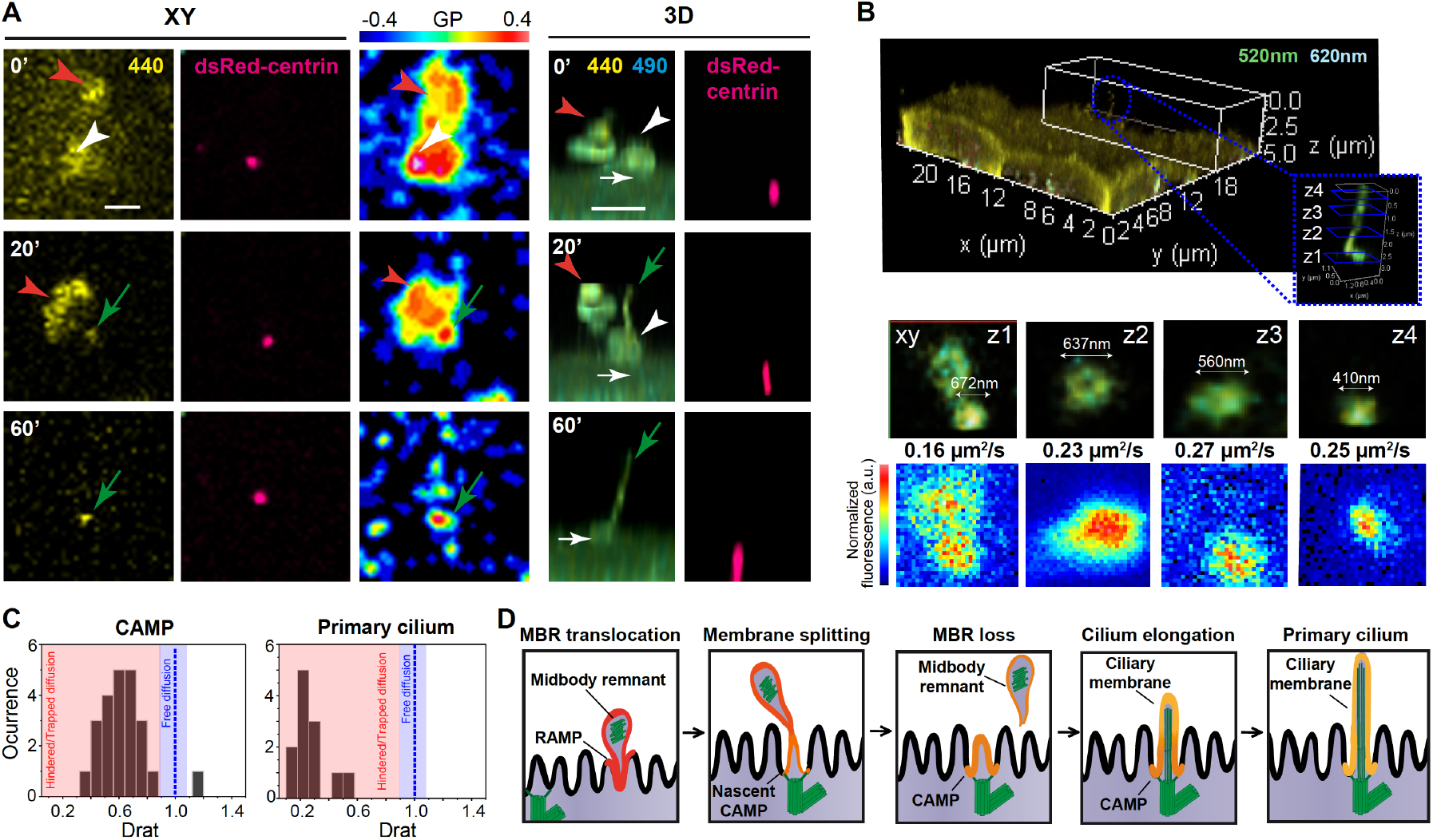
The membrane patch delivered by the MBR is used for ciliary membrane assembly. **(A)** Cells were stained with Laurdan and analyzed by videomicroscopy. The XY images show the emission at λ440 nm and the position of the centrosome, as visualized with dsRed-centrin, the corresponding GP image, and a 3D reconstruction of 8-10 stacks of the emission at λ440 and λ490 nm, and that of dsRed-centrin. A single XY plane at the level of the plasma membrane (0’) or at the middle of a primary cilium (20’ and 60’) are shown. The white and red arrowheads indicate the CAMP and the remainder RAMP membranes, respectively; the white and green arrows point to the centrosome and the nascent cilium, respectively. Scale bars, 5 µm for panoramic views and 2 µm for enlargements. **(B)** Cells were labeled with di-4-ANEPPDHQ. 3D view of a primary cilium (top panel). Merged λ560 and λ620 nm emissions are shown in green and cyan, respectively. The XY optical sections (middle panels) correspond to the z1-4 ciliary planes indicated in the inset (top panel). The diffusion map obtained by RICS analysis with indication of the mean value of the diffusion coefficient is also shown (bottom panels). **(C)** The histograms show the Drat ratio at CAMP and at the primary cilium obtained by STED-RICS analysis. Note that only hindered/trapped molecules are observed at the primary cilium. Three independent experiments, 27-28 regions of interest were analyzed. **(D)** Schematic of the proposed model for the role of the MBR in the biogenesis of the ciliary membrane. The MBR of polarized epithelial cells has a patch of condensed membranes (RAMP). The RAMP accompanies the MBR on its journey to meet the centrosome at the center of the apical membrane. Then, the RAMP delivers condensed membranes and form a new patch (CAMP) at the centrosome that is subsequently used to build the ciliary membrane.

Consistent with the CAMP being the precursor of the ciliary membrane, RICS and STED-RICS analyses showed that lipid mobility is highly restricted in both the CAMP and the primary cilium. The diffusion coefficient and the lipid lateral diffusion of the ciliary membrane were within the same range, although slightly smaller than those of the CAMP (Fig. 4B, C and Fig. S3B). This slight difference might be due to the attachment of the ciliary membrane to the axoneme through protein complexes, such as IFT trains associated with molecular motors, that might further hinder the lateral movement of lipids in the ciliary membrane. The analysis of the lipid diffusion along the z-axis of CAMPs and cilia showed that the diffusion coefficient did not change at the different planes in which lipid diffusion was measured (Fig. S3B), indicating continuity along the membrane surrounding each of these structures.

To confirm that RAMP membranes are precursors of the ciliary membrane, we photobleached RAMPs and recovered fluorescence after 3 h by re-staining with di-4-ANEPPDHQ. Whereas RAMPs were relabeled in non-ciliated cells (Fig. 2G), primary cilia were labeled in the cells that ciliated during the procedure (Fig. S3C, D). This result implies that the bleached lipids originally present in the RAMPs are transferred to form the ciliary membrane. The RAMP membranes contain machinery for primary cilium assembly, such as the IFT20, IFT81 and IFT88 subunits of the intraflagellar transport complexes (Bernabe-Rubio, et al., 2016). Therefore, it is plausible that, in addition to lipids for building the ciliary membrane, the RAMP provides the centrosome with machinery for ciliary growth.

It has been suggested that the cilium’s evolutionary origin is specialized membrane patch of the plasma membrane that recruited important sensory receptors. During evolution, the primitive patch progresses by acquiring microtubules to make it protrude for increased environmental exposure of sensory membranes. A primeval machinery derived from a coatomer-like progenitor established vectorial transport to the membrane patch (van Dam, et al., 2013). Later on, the scaffold microtubules and the patch became the axoneme and the ciliary membrane patch, respectively, and the primitive machinery evolved to the intraflagellar transport machinery (Bloodgood, 2012; Quarmby and Leroux, 2010; Jékely and Arendt, 2006). Our results showing a specialized membrane patch that is delivered by the MBR to form the ciliary membrane is somehow reminiscent of the initial part of the proposed evolutionary process. It is therefore plausible that the remnant of an ancient cytokinetic intercellular bridge constitutes a cue at the cell surface that could be associated with the evolutionary origin of the ancestral membrane patch.

Renal cystic diseases are the most common of the many abnormalities associated with ciliary dysfunction (Hildebrandt and Otto, 2005), making research on the primary cilium of renal epithelial cells particularly important. Our study identifies the source of the ciliary membrane of cells, such as that of renal epithelia, whose primary cilia are assembled entirely at the plasma membrane by revealing that the ciliary membrane arises from a patch of condensed membranes that the MBR delivers to the centrosome (Fig. 4D). These findings reveal the mechanism of biogenesis of the ciliary membrane of polarized epithelial cells and provide novel insights into how post-mitotic midbodies influence cell function.

## Materials and Methods

### Reagents

Laurdan (6-dodecanoyl-2-dimethylaminonaphthalene; product D250) and di-4-ANEPPDHQ (product D36802) were purchased from ThermoFisher. Methyl-β-cyclodextrin (product C4555) and cholesterol oxidase (product C8649) were purchased from Sigma-Aldrich.

### Cell culture

Epithelial canine MDCK II (ATCC CRL2936) and IMCD3 (ATCC CRL2123) cells were grown in MEM and DMEM/F12, respectively, supplemented with 5% fetal bovine serum (Sigma-Aldrich) at 37°C in an atmosphere of 5% CO_2_.

### DNA constructs and transfection

The DNA construct expressing dsRed-centrin2 (Tanaka, et al., 2004) (Addgene plasmid 29523) was a generous gift from Dr. J Gleeson. pEGFP-C1-MKLP1 (Addgene plasmid 70145) (Douglas, et al., 2010), which was a kind gift from Dr. M. Mishima, was used to obtain mCherry-C1-MKLP1 by standard techniques. The construct expressing mCherry-α-tubulin was from Clontech. Cells were transfected with Lipofectamine 2000 (ThermoFisher) following the manufacturer’s recommendations. Stably transfected cells were generated by transfection and selection with 1 mg/ml G-418 (Santa Cruz, CA). The resulting clones were screened under a fluorescence microscope.

### Laurdan and di-4-ANEPPDHQ staining, fluorescence microscopy and image analysis

Live cells grown on 35-mm diameter glass-bottom dishes (Ibidi-GmbH, Gräfelfing, Germany) were stained with 40 µM Laurdan for 30 min at 37°C or 5 µM di-4-ANEPPDHQ just before imaging. After excitation at λ405 nm (Gutowska-Owsiak, et al., 2018; Sezgin, et al., 2015), Laurdan intensity images were simultaneously recorded with emission in the ranges of λ420–460 nm and λ470–510 nm, which were referred to as λ440 and λ490 nm, respectively. For experiments with di-4-ANEPPDHQ, cells were excited at λ488 nm and the emission in the ranges of λ500-580 nm and λ620-750 nm was collected. These ranges were referred as λ560 nm and λ620 nm, respectively. For quantitative analysis of lipid order, a TCS SP8 inverted confocal microscope (Leica, Germany) equipped with a 405-nm laser and an HC PL APO CS2 lambda blue 63×/1.2W were used. The relative sensitivity of the two channels was determined with 0.5 mM Laurdan, and the G-factor was calculated as described (Owen, et al., 2011). For time-lapse fluorescence microscopy, cells were plated onto 35-mm glass-bottom dishes, and maintained at 37°C in MEM without phenol red containing 40 µM Laurdan and 0.25% fetal bovine serum during the recording. For quantitative analysis, cells were filmed with either the Leica microscope or a Nikon A1R confocal microscope with a 60× water objective with a numerical aperture of 1.2. For 3D reconstruction, we used NIS-Elements microscope imaging software (Nikon, Tokyo, Japan). Brightness and contrast were optimized with ImageJ (https://imagej.nih.gov/ij/) and Photoshop software (Adobe). To obtain a clearer signal in the GP images, a 2×2 pixel median smoothing algorithm was applied. For image analysis, the intensity of the emission at λ420–460 nm and λ470–510 nm was simultaneously recorded and used to calculate the pixel-per-pixel GP values according to the equation: GP=(I_440_–I_490_)/(I_440_+I_490_). 15–25 circular areas per distribution pattern, corresponding to apical plasma membrane, peripheral and central remnants, and primary cilia, were analyzed to calculate GP values, which were confirmed using SIM FCS 4 (G-SOFT Inc., Champaign, IL). GP values were represented in normalized frequency histograms and were non-linearly fitted to two Gaussian distributions using OriginPro (OriginLab, Northampton, MA). We considered acceptable only those fits for which the chi-square test gave values of p> 0.95. The area under the peaks and the maximum peak fit values were used to characterize the relative percentage of condensed membranes (coverage). Experiments were performed independently at least three times. The images shown are representative of the experiments used for quantification.

### Analysis of lipid diffusion by raster image correlation spectroscopy (RICS)

RICS was analyzed using SimFCS 4 software (G-SOFT Inc.) (Garcia and Bernardino de la Serna, 2018; Rossow, et al., 2010). To characterize the point spread function, 200 frames of freely diffusing recombinant EGFP (ranging 20-100 nM) were continuously collected in order to determine the intensity fluctuations of EGFP as a function of time. The waist of the point spread function was then adjusted by measuring the autocorrelation function of the EGFP solution with a fixed diffusion rate of 90 µm^2^/s (Garcia and Bernardino de la Serna, 2018; Rossow, et al., 2010; Digman, et al., 2005). RICS images series (256×256 pixels) were taken using either an 8, 4 or 2 μs dwell time. Subsequent analysis showed no significant difference due to different pixel dwell time. RICS analysis was performed on images after carefully observing the average intensity trace of the complete set of frames. This was done in order to identify possible photobleaching effects and to avoid potential errors in quantifying the diffusion coefficient. A dataset was discarded when the investigated set of frames showed an average intensity trace that abruptly decayed at the beginning of the collection, or when the investigated event did not show a clear disassembly process. This strategy allowed us to avoid using detrending algorithms. The RICS analysis employed a moving average (background subtraction) of 10 to discount possible artifacts arising from cellular motion and the passage slow-moving particles; smaller moving-average values were examined but revealed no difference in molecular mobility. The 2D autocorrelation map was then fitted to obtain a surface map, using the previously characterized waist value from the point spread function and the appropriate acquisition values for line time and pixel time. For different regions of interest were analyzed within the same cell, the corresponding region was drawn employing a square region of 32×32 pixels.

### Line interleaved excitation and stimulated emission depletion raster image correlation spectroscopy

Line interleaved excitation and stimulated emission depletion RICS (LIE-STED-RICS) is a bespoke implementation from a recently published line scanning STED-FCS technique, where the lipid diffusion at the microscale and nanoscale resolution were measured simultaneously (Schneider, et al., 2018). This technique allows quantification of the spatiotemporal nanoscale lipid diffusion modes and direct visualization of the macroscopic changes of membrane condensation over time. Analysis was performed with SimFCS 4 software (G-SOFT Inc.), in a similar fashion as described above, with the difference that we recorded two channels simultaneously and thereby obtained the diffusion coefficient at the microscale (D) and nanoscale (D_STED_) resolution from the RICS 2D autocorrelation fitting. To characterize the point spread function, 200 frames of freely diffusing recombinant EGFP (20 nM) were continuously collected in order to determine the intensity fluctuations of EGFP as a function of time, with variable STED laser powers. RICS images series (256×256 pixels) were taken using either an 8, 4 or 2 μs dwell time and with a 40-nm pixel size. As previously described, the 2D autocorrelation map was then fitted to obtain a surface map using the previously characterized waist value from the point spread function and the appropriate acquisition values for line time and pixel time. For different regions of interest analyses within the same cell, the corresponding region was drawn employing a square region of 32×32 pixels. All LIE-STED-RICS data were acquired with a LEICA SP8 gSTED (Leica Microsystems, Germany). For LIE-STED-RICS, we employed the alternating line scan in sequential mode, where the first line and second lines were acquired in confocal mode and STED modes, respectively. The diffusion coefficient ratios (Drat) between the diffusion coefficient values obtained from the analysis of the RICS in STED and in confocal mode (Drat=D_STED_/D) was graphed as a function of the number of events (occurrence) employing OriginPro (OriginLab, USA).

### Midbody removal

Midbody remnants were removed from cells by take-up by suction pressure as described previously with minor modifications (Bernabé-Rubio, et al., 2017; Bernabe-Rubio, et al., 2016). In brief, a heterogeneous population of MDCK cells stably expressing GFP-tubulin mixed with non-transfected cells, was grown for 4 days and stained with Laurdan. As a control, the same procedure was applied to cells in a zone of the plasma membrane distant from the MBR. Cells were re-stained with Laurdan 24 h later and analyzed for the presence of central condensed membrane patches. Cells with and without membrane patches were quantified in control cells and in cells whose MBR was removed.

### FRAP analysis

MDCK cells expressing cherry-MKLP1were stained with 5 µM di-4-ANEPPDHQ for 5 min. The di-4-ANEPPDHQ-containing medium was then washed out with PBS and replaced with normal medium. FRAP was performed in a Leica SP8 with a pulsed white light laser as an excitation source (NKT, Denmark) at a pulse frequency of 80 Mhz. In order to photobleach the selected region of interest, more than one single excitation line was chosen, as the white light laser is a much less powerful laser source than an argon laser or solid-state lasers, which are normally used to perform this technique. Lipid fluorescence was analyzed with the red laser line 555 at 5% intensity, and photobleaching of RAMPs was performed with laser lines 405, 488 and 555 at 100%. Laser power and laser line intensities were kept constant across all sample groups. To analyze the maintenance of condensed membrane structures or the appearance of a primary cilium, cells were re-stained with fresh di-4-ANEPPDHQ by carefully supplementing the medium with this compound under the microscope. For FRAP analysis, the RAMP and the plasma membrane at a region distant from the RAMP were photobleached using the same laser conditions. The FRAP bleach pulse was set to happen after 5 frames at a circular region of interest that was the same size in all the experimental repetitions. ImageJ was used to construct an intensity profile of the region of interest before and during 220 seconds after photobleaching. FRAP values were calculated as the fluorescence intensity of the photobleached region relative to the average intensity of this region before photobleaching. These values were quantified for 10 photobleached RAMPs and 10 photobleached plasma membrane regions of interest.

### Statistics

Statistical analyses were performed using GraphPad Prism 7. Results are presented as the mean ± standard derivation (SD) or standard error of the mean (SEM), as is indicated in the figure legends. Sample size and number of repeated experiments are stated in the legends. Statistically significant differences between pairs of data sets were analyzed using the two-tailed paired Student’s *t*-test. P values are shown in the figures. All experiments were performed at least three times and gave similar results.

## ACKNOWLEDGEMENTS

The expert technical advice of the Optical and Confocal Microscopy Facility is gratefully acknowledged. We thank Dr Phil Mason for revising the English language of the manuscript. This work was supported by grants (BFU2015-67266-R and PGC2018-095643-B-I00) to MAA from the Spanish Ministerio de Ciencia, Innovación y Universidades/Fondo Europeo de Desarrollo Regional (MICIU/FEDER) and by the financial support of the Central Laser Facility (Science and Technology Facilities Council, Harwell, UK) that enabled us to use in-house confocal and STED equipment under application number 16230026. EG and JBS acknowledge funding from a Marie Curie Career Integration Grant (NanodynacTCELLvation; PCIG13-GA-2013-618914). A contract from the MICIU to MB-R is also acknowledged. MB-F was supported by a fellowship from La Caixa PhD program. The authors declare no competing financial interests.

## Abbreviations used

CAMP: centrosome-associated membrane patch
FRAP: fluorescence recovery after photobleaching
GP: generalized polarization
IMCD3: inner medullary collecting duct 3
MBR: midbody remnant
MDCK: Madin-Darby canine kidney cells
MKLP1: mitotic kinesin-like protein 1
RAMP: remnant-associated membrane patch
RICS: raster image correlation spectroscopy
STED: stimulated emission depletion

---

**Figure S1.**
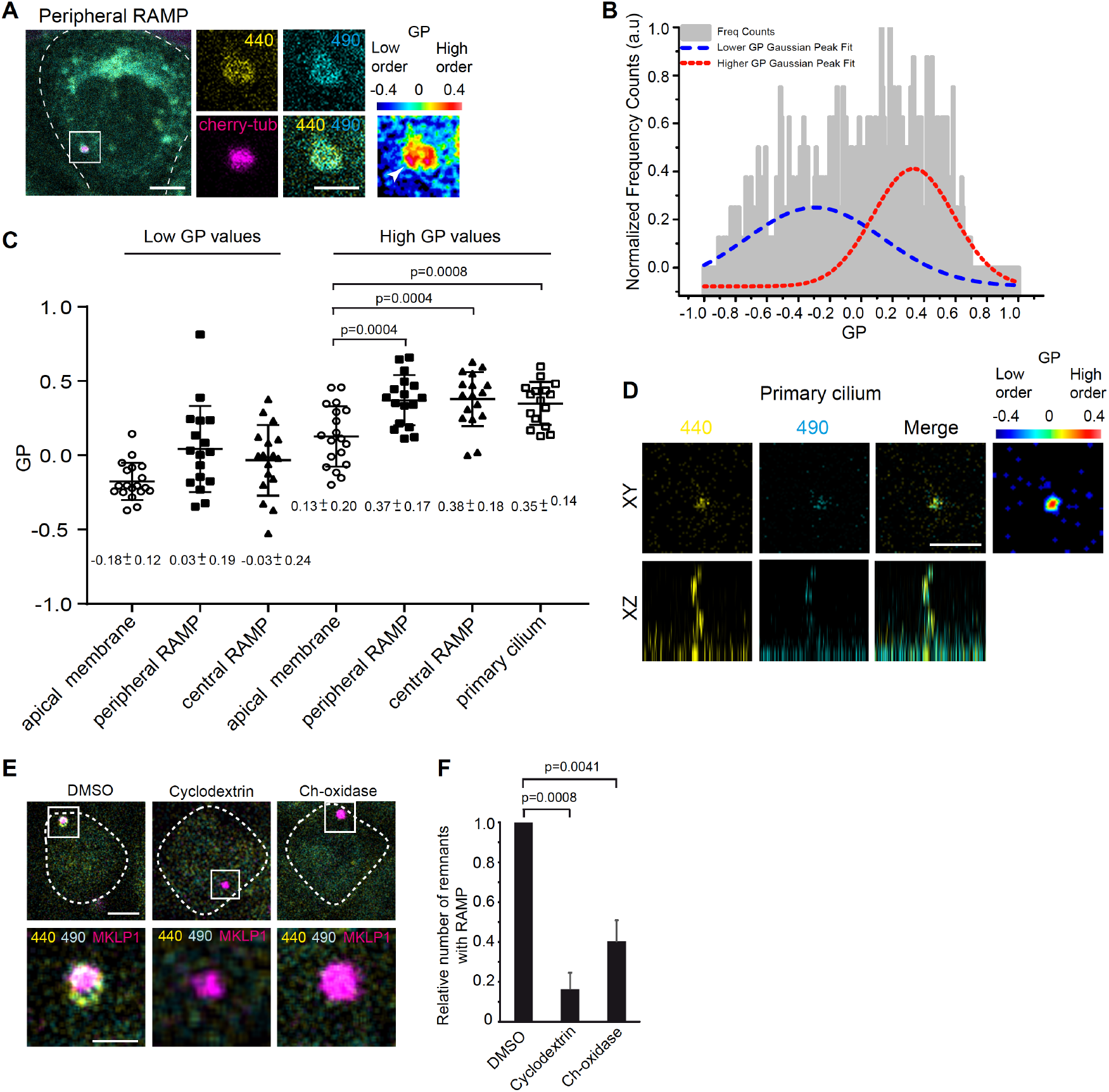
A patch of condensed membranes is associated with the MBR of IMCD3 cells. **(A)** IMCD3 cells expressing mCherry-α-tubulin were stained with Laurdan. A representative example of a cell with a peripheral MBR is shown. The enlargement of the boxed region shows the emission at λ440 and λ490 nm at the MBR area, the corresponding merge, and the localization of mCherry-α-tubulin, which was used to visualize the position of the MBR. The corresponding GP image is shown on the right. The white arrowhead indicates the MBR zone. **(B)** Calculation of GP values and coverage of membranes of a representative peripheral RAMP. The histogram shows two distinct populations of GP values. To reveal the GP of each population, we fitted two Gaussian distributions to the experimental data. The fit yielded low (blue) and highly (red) condensed membrane populations that were used to calculate the corresponding GP values. The area under each curve was used to calculate the coverage of each population. **(C)** The histogram shows the GP ± SD value corresponding to the peripheral and central RAMPs, to random zones in the apical membrane, and to cross-sections through the middle region of cilia (17-19 structures were examined in each case; three independent experiments were performed. (**D)** XY (top panels) and XZ (bottom panels) views of a representative primary cilium of IMCD3 cells. The images show the emission at λ440 and λ490 nm. The corresponding GP image is shown on the right. The GP images were colored, as indicated by the color scales. **(E, F)** MDCK cells stably expressing mCherry-MKLP1 were stained with Laurdan. Cells were then incubated with DMSO (control) or with 10 mM methyl-β-cyclodextrin or 200 U/ml cholesterol oxidase, as indicated. Representative examples of a cell with a peripheral MBR (boxed) are shown (E). The enlargement of the boxed area shows the emission at λ440 and λ490 nm in the MBR area, and that of cherry-MKLP1. Dashed lines indicate the cell contour. Scale bars, 5 µm for panoramic views and 2 µm for enlargements. (F) Relative number ± SEM of MBRs with a RAMP (25-47 MBRs were analyzed; three independent experiments were performed).

**Figure S2.**
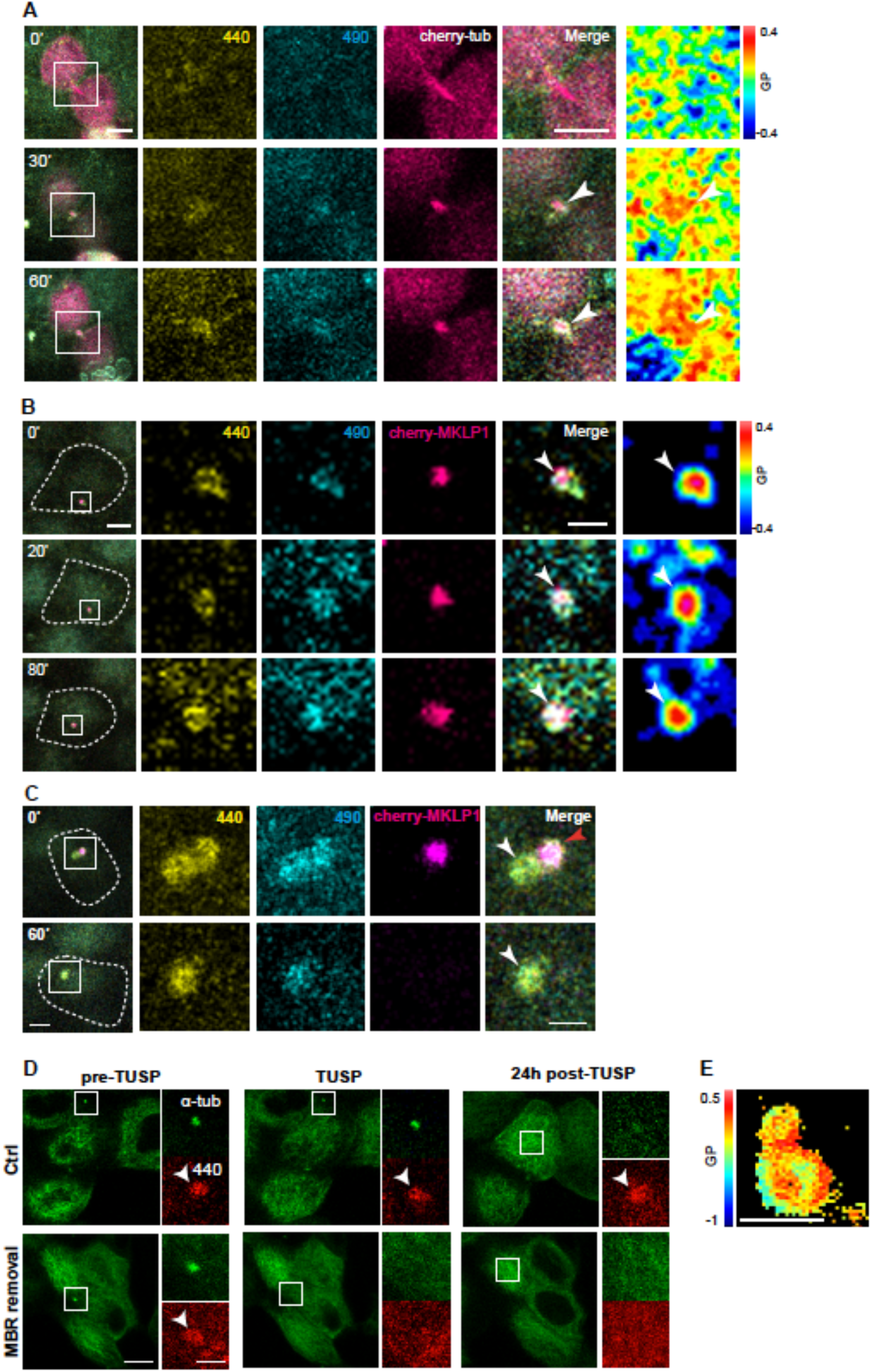
The RAMP forms soon after cytokinesis and migrates with the MBR towards the centrosome. **(A-C)** MDCK cells expressing mCherry-tubulin (A) or mCherry-MKLP1 (B,C) were stained with Laurdan and analyzed by videomicroscopy. The enlargements show the emission at λ440 and λ490 nm, and that of the exogenous proteins in the boxed area. (A) The RAMP is evident soon after cytokinesis. Since the intercellular bridge forms at the plasma membrane plane, plasma membrane staining is also visualized. (B) The RAMP moves with the MBR to a central position in the apical membrane. Corresponding GP images are also shown in (A-B). (C) After separation of the MBR from the CAMP, the MBR disappears. The white and the red arrowheads point to the RAMP and MBR, respectively. (**D**) Representative example of an experiment in which take-up by suction pressure (TUSP) was used to physically remove the MBR from the surface of cells that express GFP-tubulin to visualize the MBR. The formation of a CAMP was monitored 24 h later. As a control (Ctrl) the same procedure was applied to cells in a zone of the plasma membrane distant from an MBR. The enlargements correspond to the boxed area initially containing the MBR (pre-TUSP and TUSP) or to the center of the apical membrane (24 h post-TUSP). The white arrowhead points to the RAMP. **(E)** Cells were stained with di-4-ANEPPDHQ. The GP image obtained from the deconvolved STED image of a RAMP forming a CAMP is shown. Note the accumulation of highly condensed membranes at the interface of both structures. The dashed lines indicate the cell contour. Scale bars, 5 µm for panoramic views and 2 µm for the enlargements.

**Figure S3.**
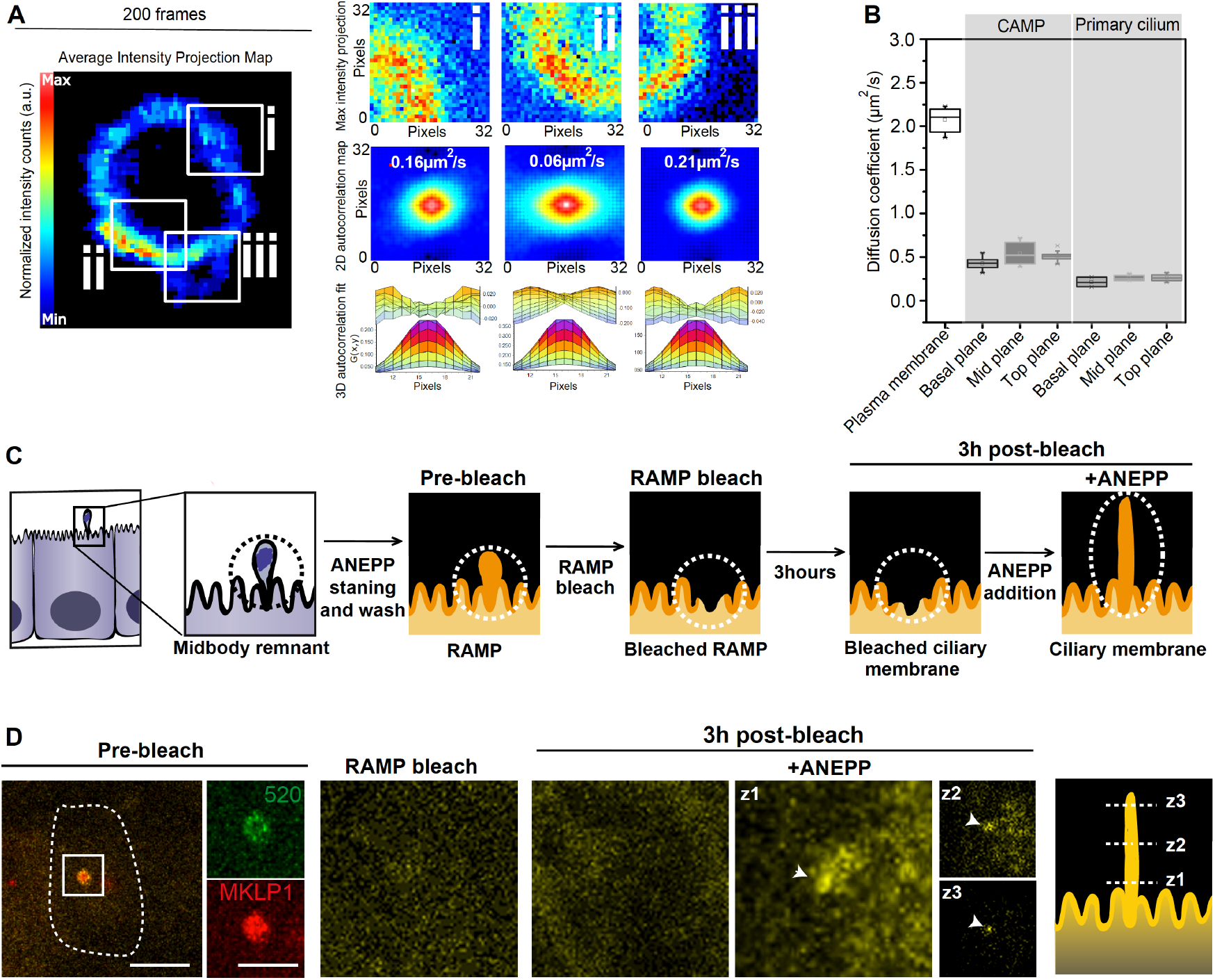
The CAMP and the ciliary membrane have similar diffusion coefficients. **(A)** Cells were labeled with di-4-ANEPPDHQ. The diffusion coefficient at RAMPs (i), at the area of CAMP emergence (ii), and at CAMPs (iii) (left panel, boxed regions) was determined by RICS analysis. Right panels show: representative maximum intensity projection maps (top panels), 2D correlation maps and diffusion coefficient (middle panels), and fit to autocorrelation function (bottom panels. For scale purposes, the pixel size is 80.4 nm and the full frame is 256×256 pixels. **(B)** RICS was used to analyze the diffusion coefficient at the apical membrane and at different transversal sections (basal, middle, top) of CAMPs and primary cilia. Note that the diffusion coefficients at CAMPs and primary cilia are in the same range in all the sections and are very different from that of the apical membrane. **(C, D)** Schematic of the FRAP experiment. RAMPs from cells expressing cherry-MKLP1 were stained with di-4-ANEPPDHQ (C). The RAMP zone (boxed) was photobleached and the cells were re-stained 3 h later. The images show a cell that has ciliated during the 3-h period (D). The RAMP was lost since it did not re-stain but, instead, the emerging cilium was labeled. The dashed line indicates the cell contour. Arrowheads indicate the position of the newly formed cilium. z1, z2, and z3 correspond to different planes along the length of the cilium. Enlargements of the boxed region are shown. Scale bars, 5 µm for panoramic views and 2 µm for enlargements.

## References

Arai, Y., Sampaio, J.L., Wilsch-Bräuninger, M., Ettinger, A.W., Haffner, C., and Huttner, W.B. (2015). Lipidome of midbody released from neural stem and progenitor cells during mammalian cortical neurogenesis. Front. Cell. Neurosci. 9, 325.

Atilla-Gokcumen, G.E., Muro, E., Relat-Goberna, J., Sasse, S., Bedigian, A., Coughlin, M.L., Garcia-Manyes, S., and Eggert, U.S. (2014). Dividing cells regulate their lipid composition and localization. Cell 156, 428–439.

Bernabe-Rubio, M., Andrés, G., Casares-Arias, J., Fernández-Barrera, J., Rangel, L., Reglero-Real, N., Gershlick, D.C., Fernández, J.J., Millán, J., Correas, I., et al. (2016). Novel role for the midbody in primary ciliogenesis by polarized epithelial cells. J. Cell Biol. 214, 259–273.

Bernabé-Rubio, M., Gershlick, D.C., and Alonso, M.A. (2017). Physical removal of the midbody remnant from polarised epithelial cells using take-up by suction pressure (TUSP). Bio-protocol 7, e2244.

Bernardino de la Serna, J., Schütz, G.J., Eggeling, C., and Cebecauer, M. (2016). There is no simple model of the plasma membrane organization. Front. Cell Dev. Biol. 4.

Bloodgood, R.A. (2012). The future of ciliary and flagellar membrane research. Mol. Biol. Cell 23, 2407–2411.

Braun, D.A., and Hildebrandt, F. (2017). Ciliopathies. Cold Spring Harb. Perspect. Biol. 9:a028191.

Chen, C.-T., Ettinger, A.W., Huttner, W.B., and Doxsey, S.J. (2012). Resurrecting remnants: the lives of post-mitotic midbodies. Trends Cell Biol. 23, 118–128.

Digman, M.A., Brown, C.M., Sengupta, P., Wiseman, P.W., Horwitz, A.R., and Gratton, E. (2005). Measuring fast dynamics in solutions and cells with a laser scanning microscope. Biophys. J. 89, 1317–1327.

Dionne, L.K., Wang, X.-J., and Prekeris, R. (2015). Midbody: from cellular junk to regulator of cell polarity and cell fate. Curr. Opin. Cell Biol. 35, 51–58.

Douglas, M.E., Davies, T., Joseph, N., and Mishima, M. (2010). Aurora B and 14-3-3 coordinately regulate clustering of centralspindlin during cytokinesis. Curr. Biol. 20, 927–933.

Elia, N., Sougrat, R., Spurlin, T.A., Hurley, J.H., and Lippincott-Schwartz, J. (2011). Dynamics of endosomal sorting complex required for transport (ESCRT) machinery during cytokinesis and its role in abscission. Proc. Nat. Acad. Sci. USA 108, 4846–4851.

Ettinger, A.W., Wilsch-Brauninger, M., Marzesco, A.-M., Bickle, M., Lohmann, A., Maliga, Z., Karbanova, J., Corbeil, D., Hyman, A.A., and Huttner, W.B. (2011). Proliferating versus differentiating stem and cancer cells exhibit distinct midbody-release behaviour. Nat. Commun. 2, 503.

Farnoud, A.M., Toledo, A.M., Konopka, J.B., Del Poeta, M., London, E., and Kenworthy, A.K. (2015). Raft-Like Membrane Domains in Pathogenic Microorganisms. In Current Topics in Membranes, Academic Press), pp. 233–268.

Garcia, E., and Bernardino de la Serna, J. (2018). Dissecting single-cell molecular spatiotemporal mobility and clustering at focal adhesions in polarised cells by fluorescence fluctuation spectroscopy methods. Methods 140–141, 85–96.

Gaus, K., Gratton, E., Kable, E.P.W., Jones, A.S., Gelissen, I., Kritharides, L., and Jessup, W. (2003). Visualizing lipid structure and raft domains in living cells with two-photon microscopy. Proc. Nat. Acad. Sci. USA 100, 15554–15559.

Gerdes, J.M., Davis, E.E., and Katsanis, N. (2009). The vertebrate primary cilium in development, homeostasis, and disease. Cell 137, 32–45.

Green, R.A., Paluch, E., and Oegema, K. (2012). Cytokinesis in animal cells. Annu. Rev. Cell. Dev. Biol. 28, 29–58.

Gutowska-Owsiak, D., Bernardino de La Serna, J., Fritzsche, M., Naeem, A., Podobas, E.I., Leeming, M., Colin-York, H., O’Shaughnessy, R., Eggeling, C., and Ogg, G.S. (2018). Orchestrated control of filaggrin-actin scaffolds underpins cornification. Cell Death Dis. 9.

Hedde, P.N., Dörlich, R., Blomley, R., Gradl, D., Oppong, E., Cato, A.C.B., and Nienhaus, G.U. (2013). Stimulated emission depletion-based raster image correlation spectroscopy reveals biomolecular dynamics in live cells. Nat. Commun. 4, 2093.

Hildebrandt, F., and Otto, E. (2005). Cilia and centrosomes: a unifying pathogenic concept for cystic kidney disease? Nat. Rev. Genet. 6, 928–940.

Ishikawa, H., and Marshall, W.F. (2011). Ciliogenesis: building the cell’s antenna. Nat. Rev. Mol. Cell. Biol. 12, 222–234.

Jékely, G., and Arendt, D. (2006). Evolution of intraflagellar transport from coated vesicles and autogenous origin of the eukaryotic cilium. BioEssays 28, 191–198.

Kuo, T.-C., Chen, C.-T., Baron, D., Onder, T.T., Loewer, S., Almeida, S., Weismann, C., Xu, P., Houghton, J.-M., Gao, F.-B., et al. (2011). Midbody accumulation through evasion of autophagy contributes to cellular reprogramming and tumorigenicity. Nat. Cell Biol. 13, 1214–1223.

Li, D., Mangan, A., Cicchini, L., Margolis, B., and Prekeris, R. (2014). FIP5 phosphorylation during mitosis regulates apical trafficking and lumenogenesis. EMBO Rep. 15, 428–437.

Lingwood, D., and Simons, K. (2010). Lipid rafts as a membrane-organizing principle. Science 327, 46–50.

Molla-Herman, A., Ghossoub, R., Blisnick, T., Meunier, A., Serres, C., Silbermann, F., Emmerson, C., Romeo, K., Bourdoncle, P., Schmitt, A., et al. (2010). The ciliary pocket: an endocytic membrane domain at the base of primary and motile cilia. J. Cell Sci. 123, 1785–1795.

Owen, D.M., Rentero, C., Magenau, A., Abu-Siniyeh, A., and Gaus, K. (2011). Quantitative imaging of membrane lipid order in cells and organisms. Nat. Protoc. 7, 24–35.

Parasassi, T., Di Stefano, M., Loiero, M., Ravagnan, G., and Gratton, E. (1994). Cholesterol modifies water concentration and dynamics in phospholipid bilayers: a fluorescence study using Laurdan probe. Biophys. J. 66, 763–768.

Pollarolo, G., Schulz, J.G., Munck, S., and Dotti, C.G. (2011). Cytokinesis remnants define first neuronal asymmetry in vivo. Nat. Neurosci. 14, 1525–1533.

Quarmby, L.M., and Leroux, M.R. (2010). Sensorium: the original raison d’être of the motile cilium? J. Mol. Cell Biol. 2, 65–67.

Reales, E., Bernabé-Rubio, M., Casares-Arias, J., Rentero, C., Fernández-Barrera, J., Rangel, L., Correas, I., Enrich, C., Andrés, G., and Alonso, M.A. (2015). The MAL protein is crucial for proper membrane condensation at the ciliary base, which is required for primary cilium elongation. J. Cell Sci. 128, 2261–2270.

Reiter, J.F., and Leroux, M.R. (2017). Genes and molecular pathways underpinning ciliopathies. Nat. Rev. Mol. Cell Biol. 18, 533–547.

Rodriguez-Boulan, E., Kreitzer, G., and Musch, A. (2005). Organization of vesicular trafficking in epithelia. Nat. Rev. Mol. Cell. Biol. 6, 233–247.

Rossow, M.J., Sasaki, J.M., Digman, M.A., and Gratton, E. (2010). Raster image correlation spectroscopy in live cells. Nat. Protocol. 5, 1761–1774.

Schneider, F., Waithe, D., Galiani, S., Bernardino de la Serna, J., Sezgin, E., and Eggeling, C. (2018). Nanoscale spatiotemporal diffusion modes measured by simultaneous confocal and stimulated emission depletion nanoscopy imaging. Nano letters 18, 4233–4240.

Sezgin, E., Levental, I., Mayor, S., and Eggeling, C. (2017). The mystery of membrane organization: composition, regulation and roles of lipid rafts. Nat. Rev. Mol. Cell. Biol. 18, 361–374.

Sezgin, E., Waithe, D., Bernardino de la Serna, J., and Eggeling, C. (2015). Spectral imaging to measure heterogeneity in membrane lipid packing. Chemphyschem. 16, 1387–1394.

Simons, K., and Gerl, M.J. (2010). Revitalizing membrane rafts: new tools and insights. Nat. Rev.Mol. Cell Biol. 11, 688.

Simons, K., and Sampaio, J.L. (2011). Membrane oganization and lipid rafts. Cold Spring Harb. Perspec. Biol. 3, a004697.

Singla, V., and Reiter, J.F. (2006). The primary cilium as the cell’s antenna: signaling at a sensory organelle. Science 313, 629–633.

Sorokin, S. (1962). Centrioles and the formation of rudimentary cilia by fibroblasts and smooth muscle cells. J. Cell Biol. 15, 363–377.

Sorokin, S.P. (1968). Reconstructions of centriole formation and ciliogenesis in mammalian lungs. J. Cell Sci. 3, 207–30.

Tanaka, K.A.K., Suzuki, K.G.N., Shirai, Y.M., Shibutani, S.T., Miyahara, M.S.H., Tsuboi, H., Yahara, M., Yoshimura, A., Mayor, S., Fujiwara, T.K., et al. (2010). Membrane molecules mobile even after chemical fixation. Nat. Meth. 7, 865–866.

Tanaka, T., Serneo, F.F., Higgins, C., Gambello, M.J., Wynshaw-Boris, A., and Gleeson, J.G. (2004). Lis1 and doublecortin function with dynein to mediate coupling of the nucleus to the centrosome in neuronal migration. J. Cell Biol. 165, 709–721.

Tyler, K.M., Fridberg, A., Toriello, K.M., Olson, C.L., Cieslak, J.A., Hazlett, T.L., and Engman, D.M. (2009). Flagellar membrane localization via association with lipid rafts. J. Cell Sci. 122, 859–866.

van Dam, T.J.P., Townsend, M.J., Turk, M., Schlessinger, A., Sali, A., Field, M.C., and Huynen, M.A. (2013). Evolution of modular intraflagellar transport from a coatomer-like progenitor. Proc. Natl. Acad. Sci. USA 110, 6943–6948.

Vieira, O.V., Gaus, K., Verkade, P., Fullekrug, J., Vaz, W.L.C., and Simons, K. (2006). FAPP2, cilium formation, and compartmentalization of the apical membrane in polarized Madin-Darby canine kidney (MDCK) cells. Proc. Nat. Acad. Sci. USA 103, 18556–18561.

