## Supplementary material for "The ciliary membrane of polarized epithelial cells stems from a midbody remnant-associated membrane patch with condensed nanodomains": Movie captions

**Movie 1**. 3D reconstruction showing that the RAMP splits into two membrane patches, one of them localizing at the centrosome. MDCK cells stably expressing dsRed-centrin stained with Laurdan were used. White and red arrowheads point to the newly formed patch and the remaining of the RAMP, respectively. The arrow indicates the position of the centrosome.

**Movie 2**. 3D reconstruction of the process of primary cilium elongation. MDCK cells stably expressing dsRed-centrin stained with Laurdan were used. White and red arrowheads point to the newly formed patch and the remaining of the RAMP, respectively; the white and green arrows point to the centrosome and the nascent cilium, respectively.
